# Specific metabolic signatures of fish exposed to cyanobacterial blooms

**DOI:** 10.1101/416297

**Authors:** Benoît Sotton, Alain Paris, Séverine Le Manach, Alain Blond, Charlotte Duval, Qin Qiao, Arnaud Catherine, Audrey Combes, Valérie Pichon, Cécile Bernard, Benjamin Marie

## Abstract

With the increasing impact of the global warming, occurrences of cyanobacterial blooms in aquatic ecosystems are becoming a main ecological concern around the world. Due to their capacity to produce potential toxic metabolites, interactions between the cyanobacteria/cyanotoxin complex and the other freshwater organisms have been widely studied in the past years. Non-targeted metabolomic analyses have the powerful capacity to study a high number of metabolites at the same time and thus to understand in depth the molecular interactions between various organisms in different environmental scenario and notably during cyanobacterial blooms. In this way during summer 2015, liver metabolomes of two fish species, sampled in peri-urban lakes of the île-de-France region containing or not high concentrations of cyanobacteria, were studied. The results suggest that similar metabolome changes occur in both fish species exposed to cyanobacterial blooms compared to them not exposed. Metabolites implicated in protein synthesis, protection against ROS, steroid metabolism, cell signaling, energy storage and membrane integrity/stability have shown the most contrasted changes. Furthermore, it seems that metabolomic studies will provide new information and research perspectives in various ecological fields and notably concerning cyanobacteria/fish interactions but also a promising tool for environmental monitoring of water pollutions.

## Introduction

Direct and indirect anthropogenic disturbances affect the global functioning and stability of freshwater ecosystems. Global warming, physical alteration, habitat loss, water withdrawal, pollutions, overexploitation and the introduction of non-native species constitute the main threats affecting freshwater ecosystems and their biocenoses (Bunn, 2016; Dudgeon et al., 2006; Revenga et al., 2005). Chemical pollutions in freshwater environments have noticeably become a main concern: more than 60 000 man-made chemicals are produced and regularly used for human activities, and being potentially transferred to natural ecosystems (Hamilton et al., 2016; Schwarzenbach et al., 2010). Over the past century, transformation of natural landscapes for industrial and urban needs have led to a general increase of chemical nutrients concentrations in freshwater ecosystems, and notably of phosphorus and nitrogen species that have caused the massive eutrophication of freshwater bodies (Schwarzenbach et al., 2010). One of the main consequences of this non-natural eutrophication process is a shift in the composition of the primary producers due to the appearance of noxious autotrophic bacteria, such as cyanobacteria (Chorus and Bartram, 1999; O’Neil et al., 2012). In worldwide freshwater ecosystems, cyanobacteria are mainly present during summers, forming large surface scums generally accompanied by the presence of toxic compounds that have been pointed out to affect all trophic food web compartments (Codd et al., 2005; Ferrão-Filho & Kozlowsky-Suzuki, 2011; Sotton et al., 2014). Furthermore, the potential inducing effect of the global warming on the frequency, the intensity and the occurrence of cyanobacterial blooms have placed these phenomena as a main threat to public health and ecosystem sustainability (Brooks et al., 2016; O’Neil et al., 2012; Sukenik et al., 2015). In freshwater ecosystems, *Microcystis, Aphanizomenon, Anabaena, Cylindrospermopsis* and *Planktothrix* are the main bloom-forming cyanobacterial genera found during summers in lentic waterbodies, and all have been often highlighted to produce notorious toxic compounds, so-called cyanotoxins (Codd et al., 2017). Among them, the microcystins (MCs), a family of hepatotoxins consisting of more than 230 variants, are the most studied due to their high biological activity and their wide occurrence during freshwater cyanobacterial blooms (Catherine et al., 2017). MCs may induce to fauna inhibition of the protein phosphatases 1 (PP-1) and 2A (PP-2A) as well as occurrence of a cellular oxidative stress *via* the formation of reactive oxygen species (ROS), with different physiological responses and consequences depending on the organism and the species studied (Amado and Monserrat, 2010; Malbrouck and Kestemont, 2006). The effects of freshwater cyanobacteria and their respective cyanotoxins, notably the MCs, have been widely studied on the ichtyofauna, constituting one of the most potentially relevant indicators of environmental disturbances (Bols et al., 2001; Malbrouck & Kestemont, 2006). However, the actual knowledge dealing with cyanobacteria impairs to natural population fishes is mainly based on experimental data reporting, generally after short-term exposures using high concentrations of purified toxins, or the mechanisms involved in the accumulation-detoxification dynamics of MCs. To our opinion, there is still a lack in the deep understanding, particularly on the ichtyofauna, of the real ecotoxicological effects of natural cyanobacterial biomasses, producing at the same time a “cocktail” of potentially bioactive compounds. To an ecological point of view, the metabolome of an organism is the set of primary metabolites synthesized at a given time and thus represents its metabolic chemical picture, which can potentially be altered when the ecological and environmental stress conditions change (Bundy et al., 2008; Fiehn, 2002). Study of the metabolome regulations represents the final endpoint of the phenotypic response of an organism that allows it to counter-act the potential negative effects of stressors present in ecosystems, and thus to adapt to the variable conditions. In this way, as changes at the metabolome scale are directly influenced by transcriptome and proteome changes, metabolomic studies have become a relevant approach to describe and analyze an integrated response of an organism under specific environmental context and scenario (Bundy et al., 2008; Franzosa et al., 2015; Hultman et al., 2015; McLean, 2013). The changes in the primary metabolite concentrations help to provide valuable and useful information concerning the physiological processes involved in the homeostatic responses of the organisms encountering environmental stresses from potentially multiple origins. Nuclear magnetic resonance (NMR)-based metabolomics has been proved to be a powerful approach to address hypotheses relating to fish physiology and development or pollutant induced toxicity or diseases (Samuelsson & Larsson, 2008; Sardans et al., 2011; Viant, 2008). However, despite its high potential to better understand the molecular mechanisms implicated in the ecotoxicological responses of organisms, the investigation of new qualitative and quantitative biomarkers showing the interactions between biocenoses and their biotopes in various ecological context remains still rare (Bundy et al., 2008; Samuelsson and Larsson, 2008; Sardans et al., 2011; Viant, 2008).

Thus, as toxic cyanobacterial blooms may represent important ecotoxicological and ecological constraints in freshwater environments, it can be supposed that organisms exposed to cyanobacterial blooms exhibit characteristic metabolome signatures, compared to others non-exposed to cyanobacterial dominant conditions, as their metabolism may respond and counter-act the potential negative effects of the cyanobacteria and their cyanotoxins and thus adapt to the local environmental pressures. To date and to our knowledge, no studies have been carried out using a metabolomic approach in order to assess the specific metabolic changes that could be observed in fish exposed to cyanobacterial blooms in contrasted aquatic ecosystems. In this way, during the summer 2015, individuals of two representative fish species of freshwater lakes from the European temperate regions, the perch (*Perca fluviatilis*) and the pumpkinseed sunfish (*Lepomis gibbosus*), have been sampled in eight peri-urban lakes of the île-de-France region contrasted by their phytoplanktonic community composition (“presence” or “absence” of cyanobacterial blooms). ^1^H-NMR metabolomic analyses were performed on the fish liver in order to investigate the global metabolome responses of the two fish species exposed to distinct ecological scenario and to further identify the metabolic signatures related to these potential specific phenotypic responses.

## Materials and methods

### Lakes and physico-chemical parameters measurements

*In-situ* sampling campaigns were performed during summer 2015 in eight lakes of the île-de-France region, chosen for their different dominant phytoplankton communities already described in previous studies (Catherine et al., 2008; Maloufi et al., 2016) and notably the presence or the absence of recurrent cyanobacterial blooms. Les Galets lake (C), La Sablière lake (CM), La Courance lake (M), Saint Cucufa lake (R), Grosse Pierre lake (V), Grand Marais lake (VS), Grand Fontenay lake (F) and Triel lake (T) were thus studied and sampled with electric fishing device for capturing fish alive. In all lakes, dissolved oxygen (O_2_) concentrations, pH, temperatures and conductivity were measured in the water column using a multiparameter probe (YSI EXO2).

### Phytoplankton sampling

In every lake, sub-surface chlorophyll-*a* equivalent concentrations attributed to the four-main phytoplankton groups (Chlorophyta, Diatoms, Cyanobacteria, Cryptophyta) were measured with an *in-situ* fluorometer (Fluoroprobe II, Bbe-Moldenke, Germany) and samples of water were filtered through 1.2 μm GF/C filters (Nucleopore, Whatman) and stored at −80°C until total chlorophyll-*a* concentrations analyses using the ethanolic extraction as described by Yepremian *et al.* (2017). Samples of water were fixed in Lugol iodine solution and kept at 4°C until the identification. The estimation (%) of the abundance of the different cyanobacterial genera was performed using an Utermohl’s counting chamber and an inverted microscope as described by Catherine et al. (2016). In parallel, phytoplankton biomass was collected using an Apstein’s type phytoplankton net (20-μm mesh size) and kept at −80°C until metabolite characterization by mass spectrometry (MS) and MCs content analyses.

### Metabolite characterization of phytoplanktonic biomasses by mass spectrometry

The biomasses from the sampled lakes were freeze-dried and then sonicated in 80% methanol, centrifuged at 4°C (4,000 g; 10 min). The supernatant was transferred and acidified with formic acid and 5 μL were analyzed on an HPLC (Ultimate 3000, ThermoFisher Scientific) coupled to an electrospray ionization and quadrupole time-of-flight hybrid mass spectrometer (ESI-QqTOF, QStar® Pulsar i, Applied Biosystems®, France).

High-performance liquid chromatography (HPLC) of 5 *μ*L of each of the metabolite extracts was performed on a C_18_ column (Discovery® Bio wide pore, 15cm * 1mm, 5 μm, Sigma) at a 50 μL.min^-1^ flow rate with a gradient of acetonitrile in 0.1% formic acid (10 to 80% for 60 min). The metabolite contents were then analyzed on positive mode using information dependent acquisition (IDA), which allows switching between mass spectrometry (MS) and tandem mass spectrometry (MS/MS) experiments, as previously described (Marie et al., 2012). The data was acquired and analyzed with the Analyst QS software (Version 1.1). Peak lists were generated from MS spectra acquired between 10 and 55 min, filtering noise threshold at 2% maximal intensity and combining various charge states and related isotopic forms. Metabolite annotation was attempted according to the accurate mass of the molecules, then to their respective MS/MS fragmentation pattern with regard to an in-house database of more than 800 cyanobacterial metabolites.

### Cyanotoxins quantification

For MCs extraction, phytoplankton biomasses (50 mg dry weight) of each lake were placed in a glass tubes, extracted with 5 mL of 75% methanol and sonicated for 5 min in an ultrasonic bath. This step was performed twice. The crude extracts were centrifuged at 20,000 rpm for 15 min at 4°C. The supernatants were then collected with a Pasteur pipet and kept at −80°C until MCs analysis by ELISA, done using the microcystins (Adda-specific) Kit (Abraxis LLC) recommendations. Prior to analysis, samples were dissolved with the ELISA sample diluent to reach a methanol concentration below 5% to avoid any interactive effect and to stay in the detection range of the kit (0.1-5 μg.L^-1^) for all samples. The results were expressed in MC-LR equivalents (μg eq. MC-LR .mg^-1^ dry weight).

We also attempt to measure free and bound BMAA in the phytoplankton biomass according to the HILIC/MS-MS based methods described previously (Combes et al., 2014; Faassen et al., 2016). This method used solid-phase extraction based on mixed mode sorbent to concentrate and clean up the phytoplankton extract that contained free BMAA. After the acidic lysis of the phytoplankton biomasses, the bound fraction of BMAA was analyzed by LC/MS-MS. This quantitative method has proved to be specific and reliable in a range of concentration level from 0.25 to 1.6 ng mg^-1^ dry weight.

### Fish sampling and tissue extraction procedure for metabolomic analysis

Immature individuals of perch (*Perca fluviatilis*) and pumpkinseed sunfish (*Lepomis gibbosus*), two representative fish species of European freshwater lakes, were targeted by electric fishing (FEG 8000, EFKO, Leutkirch, Germany) performed in the riparian area of every lake. Alive fish caught were directly measured, weighed (Table S2), euthanized and then liver of each individual was shortly sampled, deep-frozen in liquid nitrogen and kept at −80°C until metabolomics analyses.

Liver metabolome extraction was carried out using the methanol/chloroform/water (ratio 2/2/1.8) method, on the basis of existing literature (Lin et al., 2007; Wu et al., 2008). Briefly, fresh frozen livers were weighed, homogenized in ice-cold methanol (8 mL per gram of tissue; AnalaR Normapur, min. 99.8 %, VWR, Pennsylvania, USA) and ice cold milliQ water (2.5 mL.g^-1^), and then vortexed for 1 min. Subsequently, ice cold chloroform (4 mL.g^-1^; Normapur, 99.3 %, VWR, Pennsylvania, USA) and milliQ water (4 mL.g^-1^) were added to extract the hydrophobic metabolites. Then, the mixture was vortexed for 1 min and incubated on ice for 10 min to obtain a complete solvent partition. The resulting supernatant was then centrifuged at 4°C for 10 min at 2,000 g, resulting in a biphasic solution. The upper polar and lower non-polar layers were carefully removed. The upper polar fraction was then transferred to 2-mL Eppendorf tubes, dried under Speed-vac device (Speed-vac Plus SC110A, Savant) and then kept at −80°C until NMR analysis. Prior to ^1^H-NMR measurement, the polar tissue extracts were dissolved in 550 μL of 0.1 M sodium phosphate buffer prepared in D_2_O (10% v/v) containing 0.25 mM sodium-3-tri-methylsilylpropionate (TMSP) as an internal standard. Finally, the resulting samples were transferred to 5-mm NMR tubes (Norell, France) and immediately analyzed by ^1^H-NMR.

### 1 H-NMR spectroscopy

All NMR data were recorded at 298 K on a 600 MHz Bruker AVANCE III HD spectrometer equipped with a 5-mm TCI CryoProbe (^1^H-^13^C-^15^N) with Z-gradient. One-dimensional ^1^H-NMR spectra were acquired using a standard Bruker noesygppr1d pulse sequence to suppress water resonance. Each spectrum consisted of 256 scans of 32768 data points with a spectral width of 7.2 kHz, a relaxation delay of 2.5 s and an acquisition time of 2.3 s.

### BATMAN metabolite quantification

The relative metabolite quantification was performed using the BATMAN (an acronym for Bayesian AuTomated Metabolite Analyser for NMR spectra) R-package (Hao et al., 2014), which deconvolutes peaks from 1-dimensional ^1^H-NMR spectra to automatically assign chemical shifts to specific metabolites from a target list and then estimate their respective concentrations. This can be achieved thanks to an implementation of a Bayesian-based modelling procedure. BATMAN uses, in a two-component joint model, resonances of every assigned proton from a list of catalogued metabolites and noisy information to finally reconstruct the empirical NMR spectrum. But, in absence of confirmatory analytical methods, we prefer to state them as candidate metabolic biomarkers sharing the same ^1^H-NMR parameters with the catalogued metabolites. Therefore, 222 metabolites were quantified from Bruker spectra files using the following parameters: i) the chemical shift regions belonging to the two following regions: 0.5 to 4.60 ppm and 5.40 to 10.0 ppm, ii) 400 burn-in iterations, iii) 200 post-burn-in iterations and iv) 5000 iterations for batman rerun. Calculations were performed on a HP Z820 workstation using two 3.30 GHz Intel Xeon^©^ CPU E5 processors and 64 Go RAM by activating 12-parallel threads processing.

### Statistical exploration of data

The mixOmics library was used to carry out the multivariate analyses. Regularized canonical correlation analysis (rCCA) is a multivariate statistical method used to assess correlations between two multivariate datasets acquired on the same individuals. Here, it was used as a factorial discriminant analysis that modeled the relationships between, the species (*Perca* and *Lepomis*), the chemical data measured in the different lakes (total Chl-*a* concentrations, Chl-*a* estimated concentrations related to cyanobacterial biomasses (BBE), MCs concentrations, O2 concentrations, pH and Conductivity), and the semi-quantitative levels of metabolites determined by the BATMAN algorithm. The *rcc()* function was used to define the canonical correlations and the canonical variates; the *network()* function was used to produce the network of interactions. Cross-validation of MANOVA results was obtained thanks to a bootstrap-based procedure by applying a random assignment of any statistical individual to a given group of treatments. In order to evaluate the effects of the experimental factors (species, chemical data and their interaction) on the relative concentrations of metabolites highlighted by multivariate analyses and the subsequent relevant networks, simple two-way ANOVAs followed by a Student-Newman-Keuls post-hoc test were performed.

## Results

### Phytoplankton, microcystins and chemical conditions of the studied lakes

During 2015 summer samplings, distinct phytoplanktonic compositions and concentrations were observed in the eight lakes targeted (Fig. S1). Among them, F, T, V and VS, exhibited remarkable high and/or dominance of cyanobacterial-specific chlorophyll-*a* concentrations reaching 106, 14.8, 135 and 48.3 μg eq. Chl *a* L^-1^, respectively (Fig. 1A). In the other lakes, chlorophyll-*a* concentrations of the total phytoplanktonic community ranged between 7.2 and 241.5 μg. Chl *a* L^-1^ (Table S1) but were mainly dominated by Chlorophyta, diatoms and/or cryptophytes (Fig. S1), major phytoplanktonic phyla, that are so far not known to produce any toxins. In these lakes, only low concentrations of cyanobacterial-specific chlorophyll-*a*, ranging from 2.9 to 10.9 μg eq. Chl *a* L^-1^, were measured (Fig. 1A). Furthermore, in the cyanobacteria-dominated lakes, different cyanobacterial genera were observed during samplings (Fig. 1B). F and T were dominated by *Planktothrix* (94% of the cyanobacteria present in the sample) and *Pseudanabaena* (71%), respectively, whereas V and VS, were both dominated by *Microcystis* (Fig 1B).

**Figure 1:**
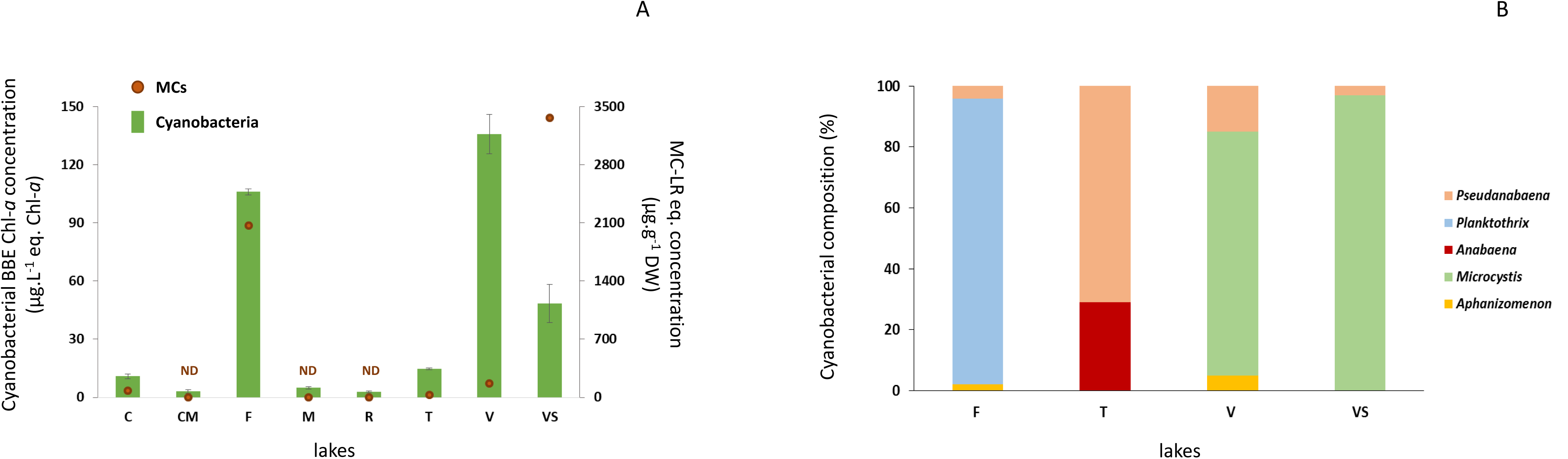
Cyanobacterial specific chlorophyll–*a* concentrations (μg.L^-1^ eq. Chl-a) and MCs concentrations (μg eq. MC-LR.g^-1^ DW) in the sampled lakes (A) and the cyanobacterial genus composition observed in the cyanobacteria-dominated lakes (B). ND = non-detected. Les Galets lake (C), La Sablière lake (CM), La Courance lake (M), Saint Cucufa lake (R), Grosse Pierre lake (V), Grand Marais lake (VS), Grand Fontenay lake (F) and Triel lake (T).

MC-LR equivalent concentrations were measured at various concentrations ranging from above the quantification limit (> 1 μg.g^-1^ DW) to 3367 μg.g^-1^ DW eq. MC-LR (Fig 1A, Table S3). The highest MCs concentrations were found in lakes dominated by cyanobacteria and particularly in F, V and VS where 2067.3 μg.g^-1^, 166.1 μg.g^-1^ and 3367 μg.g^-1^ MC-LR eq. were measured, respectively (Fig 1A, Table S3). Interestingly, V that exhibited the highest cyanobacterial concentrations (dominated by *Microcystis*) measured in our study, did not exhibit the highest MCs concentrations (3367 μg.g^-1^ MC-LR eq. in VS) illustrating the fact that MCs concentrations are not only linked to the cyanobacterial biomass measured in the lakes (Fig. 1A, Table S3). However, none of these samples presented detectable amount of the potential neurotoxin BMAA (neither in free or bound form). Nevertheless, various classes of cyanobacterial secondary metabolites together with different variants of microcystins, anabaenopeptins, aeruginosins, microginins and cyanopeptolins were detected in the dried biomasses of lakes where cyanobacteria reached high concentrations (Table S4).

### Metabolomic analysis of sampled fish

^1^H-NMR raw files were preprocessed thanks to the R BATMAN library (Hao et al., 2014) to get a relative quantification of a preselected set of metabolites replacing an analytical assignment based on different and complementary analytical methods. Thus, a confident quantification of each metabolite determined on each spectrum is given, even for very low concentrated metabolites when compared to the main ones, as the differences in their relative concentrations could have been validated by both uni- and multivariate statistical tests.

A multivariate analysis was performed by using a rCCA analysis between the matrix of the relative concentrations of the 222 metabolites (X) and the dummy matrix (Y) corresponding to the following factors: *i.e.* fish species, cyanobacterial-specific Chl-*a* and MCs concentrations, O_2_, pH, conductivity and the total Chlorophyll-*a* concentrations and their interaction using the mixOmics package in R (version 3.2.0) and a MANOVA bootstrap procedure applied to the dataset in order to highlight the significant effects of the environmental factors on the observed metabotypes (*i.e.* the specific metabolic profiles according to the environmental factors). MANOVA bootstraps reveal significant effects of species (*p* = 0), cyanobacterial-specific chl-*a* (BBEcya) concentrations (*p* < 10^−43^), MC concentrations (*p* < 10^−11^), pH (*p* < 10^−28^), O_2_ (*p* < 10^−28^), conductivity (*p* < 10^−47^), Total chlorophyll-a concentrations (*p* < 0.011) from the rCCA model. However, a clear species effect is observable through the dimension 1 (*p* < 10^−16^) with *Perca* and *Lepomis* clearly separated by the first dimension whatever the lake considered (fig. 2A). On the dimension 2, a clear cyanobacterial concentration effect is observable (*F* = 1931, *p* < 10^−16^), with fish coming from cyanobacterial dominated lakes (in green) clearly separated by this dimension whatever the fish species considered (Fig. 2A). In addition, O_2_ concentrations (*F* = 599, *p* < 10^−16^), pH (F = 867, *p* < 10^−16^) and conductivity (*F* = 419, *p* < 10^−16^) contribute significantly to the different metabotypes observed on this dimension but to a much lesser extent compared to cyanobacterial concentrations (Fig. 2A). Finally, no significant interactive effects between species and the environmental factors have been observed in our analysis suggesting that a similar response of the two species to the different environmental factors occurs.

**Figure 2:**
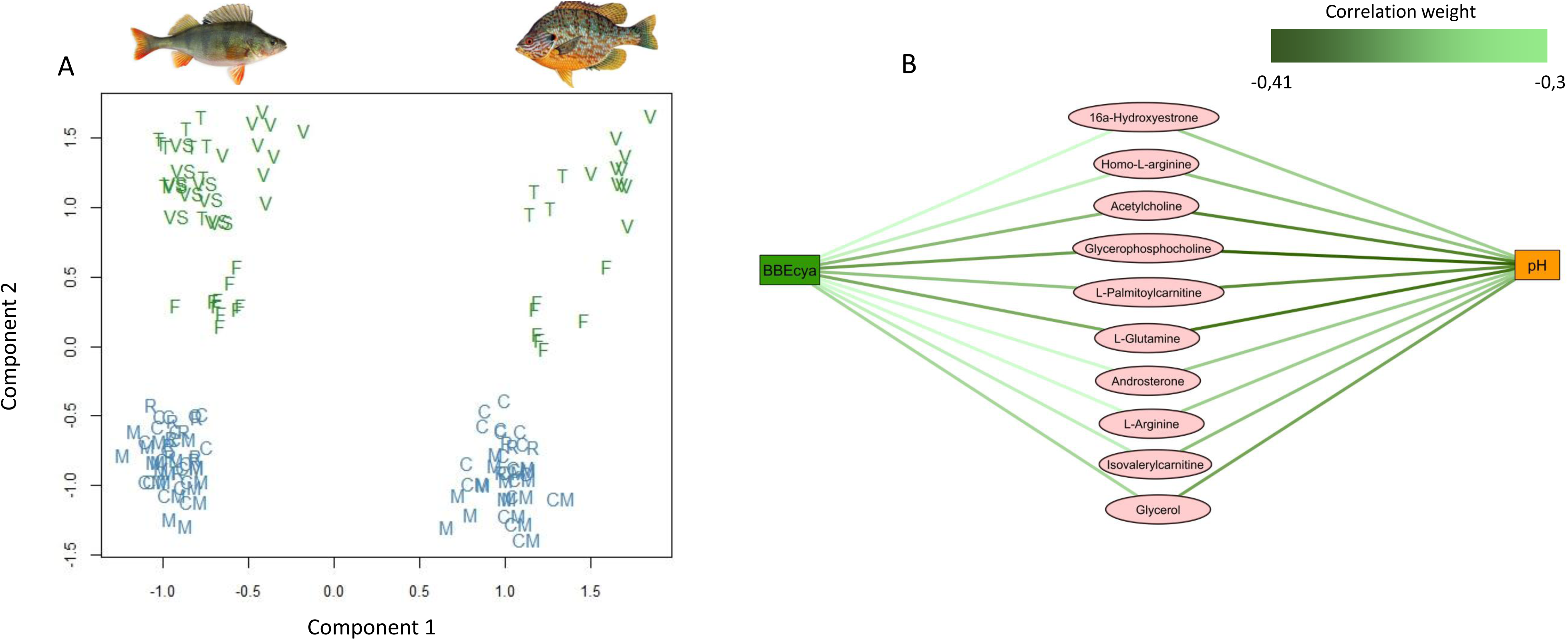
^-1^H-NMR liver metabolomes and relevance network of fish sampled in the different lakes. The individual plots of regularized canonical correlations analysis (rCCA) for dimensions 1–2 (A). Perch individuals are on the left side and pumpkinseed individuals on the right side of the graphic. Lakes are represented by their respective letters that in blue correspond to control lakes and in green to perturbated lakes. Relevance network providing from rCCA analysis on the dimension 2(B). Putative metabolites above a correlation threshold of 0.3 were kept. Green edges correspond to negative correlations with the discriminant ecological factors.

The relevance network based on the second rCCA dimension specifically highlights metabolites, above a |correlation| threshold of 0.3, discriminating the fish exposed or not to the cyanobacteria and linked to the environmental factors responsible for the discrimination (mainly BBEcya concentrations but also pH) (Fig. 2B). It appears that all these metabolites shown by this network exhibit negative correlation with both cyanobacterial concentrations and pH values. In addition, two-way ANOVAs were performed on all putative metabolites highlighted in the figure 2B (Table 1). A significant effect of the BBEcya and pH is shown for 16a-hydroxyestrone, acetylcholine, androsterone, glycerophosphocholine, isovalerylcarnitine, L-glutamine and L-palmitoylcarnitine whereas for glycerol, homo-L-arginine and L-arginine, a significant effect of the BBEcya concentrations, only, have been confirmed by two-way ANOVA analyses (Table 1). Furthermore, no interaction effect between the species and the BBEcya concentrations or pH has been observed for any metabolites suggesting similar metabolic responses, based on the metabolites highlighted by our relevance network, for these two species and for these two discriminating environmental factors. Negative correlations between relative concentrations of the metabolites and the BBEcya concentrations and the pH conditions are translated by significant lowest relative concentrations of the metabolites in the liver of fish captured in cyanobacterial dominated lakes (Figure S2). Globally, it appears that liver of fish sampled in T, F and V lakes exhibits the most significant differences in the relative concentrations of the highlighted metabolites compared to fish coming from lakes where cyanobacteria are in low or not dominant concentrations.

**Table 1:** Effects of significantly-related environmental factors on the metabolite expressions revealed by two-ways ANOVA analyses. Bold *p*-values correspond to *p*-values < 0.05.

| Metabolites | BBECya |  |  | pH |  |  |
| --- | --- | --- | --- | --- | --- | --- |
|  | *Df* | *F-value* | *p-value* | *Df* | *F-value* | *p-value* |
| 16a-Hydroxyestrone | 1 | 26.604 | **1.02e<sup>-06</sup>** | 1 | 8.295 | **0.00472** |
| Acetylcholine | 1 | 43.903 | **1.08e<sup>-09</sup>** | 1 | 5.892 | **0.01672** |
| Androsterone | 1 | 19.932 | **1.85e<sup>-05</sup>** | 1 | 10.776 | **0.00135** |
| Glycerophosphocholine | 1 | 38.040 | **1e<sup>-08</sup>** | 1 | 8.892 | **0.00348** |
| Glycerol | 1 | 29.455 | **3.1e<sup>-07</sup>** | 1 | 3.908 | 0.0504 |
| Homo-L-arginine | 1 | 32.501 | **8.95e<sup>-08</sup>** | 1 | 1.795 | 0.18287 |
| Isovalerylcarnitine | 1 | 32.028 | **1.08e<sup>-07</sup>** | 1 | 9.392 | **0.0027** |
| L-Arginine | 1 | 30.269 | **2.22e<sup>-07</sup>** | 1 | 2.228 | 0.13820 |
| L-Glutamine | 1 | 30.004 | **2.47e<sup>-07</sup>** | 1 | 14.025 | **0.000280** |
| L-Palmitoylcarnitine | 1 | 24.965 | **2.05e<sup>-06</sup>** | 1 | 12.888 | **0.000483** |

## Discussion

### Diversity of ecosystems: environmental, chemical and biological conditions

Cyanobacterial blooms are frequent phenomenon in peri-urban lakes from the île-de-France region due to the eutrophic and hyper-eutrophic states of these environments. Interestingly, these freshwater aquatic ecosystems are characterized by a wide range of environmental conditions and anthropogenic pressures that influence the phytoplankton biomasses and compositions present in these ecosystems (Catherine et al., 2010, 2016; Maloufi et al., 2016). *Microcystis* genera was largely dominant in V and VS lakes with cyanobacterial-specific chl-*a* (BBEcya) concentrations reaching 135 and 48.3 μg eq. chl-*a* L^-1^, respectively, whereas in C the BBEcya concentration did not exceed 11 μg eq. chl-*a* L^-1^ and *Microcystis* cells were co-dominant with Chlorophyta. In F, *Planktothrix* genera was found to be dominant with BBEcya concentration reaching 106 μg eq. chl-*a* L^-1^. Interestingly, in T lake, *Pseudanabaena* and *Anabaena* genera were present at low BBEcya concentrations (around 15 μg eq. chl-*a* L^-1^) compared to those measured in the other cyanobacteria dominated lakes (F, V, VS). However, even though cyanobacterial amount in T was not as high as in cyanobacterial-dominated lakes of this study (F, V and VS), the long-term presence in the T lake of noticeable amount of other contaminants, such as heavy metals, phthalates, pesticides, polychlorinated biphenyls (PCBs), estrogen hormones and polycyclic aromatic hydrocarbons (PAHs) that have been measured in both water and fish samples (Azimi and Rocher, 2016; Gaspéri et al., 2010; Teil et al., 2014), would explained why the fish present in this environment exhibit an altered metabolic profile, similar to those of the cyanobacteria-dominated lakes. In the other lakes (R, C, M and CM), diatoms or chlorophyta were dominant and cyanobacteria were present with chl-*a* concentrations never exceeding 5 μg eq. chl-*a* L^-1^.

All cyanobacterial genera encountered in these lakes are commonly observed during bloom episodes occurring in European lentic freshwaters (Padisak et al., 2016) and are susceptible to represent an ecological risk due to their capacity to produce toxic secondary metabolites, such as MCs (Bernard et al., 2017; Buratti et al., 2017; Oudra et al., 2002). The highest MCs values were measured in VS (3367 μg eq. MC-LR .g^-1^ DW) and then in F (2067.3 μg eq. MC-LR .g^-1^ DW). Furthermore, despite the higher cyanobacterial chl-*a* content observed in V compared to VS and F, lower MCs concentrations have been measured in this lake (166 μg eq. MC-LR .g^-1^ DW). This observation could be linked to the presence of both MC-producing and non-producing genotypes of *Microcystis* as it is known that several genotypes within one cyanobacterial species can co-occur during bloom episodes (Sabart et al., 2010). However, supplementary studies should be performed in order to verify this hypothesis. Nevertheless, cyanobacteria are well known to produce a wide variety of potentially toxic secondary metabolites other than MCs, and our global network analysis of the metabolite production of the cyanobacterial biomasses of the various lakes highlights that other known (cyanopeptolin, aeruginosin, microginin and anabaenopeptin) and various unknown metabolites were also present, supporting necessary in-depth investigation of the molecular diversity of the metabolite produced by phytoplankton biomasses (among which cyanobacterial biomass) in natural aquatic ecosystems. The potential ecological consequences of the production of this chemical diversity of metabolites by cyanobacteria remain largely uncovered, as some of these components are so far described to display molecular bioactivity as inhibitory effects on different proteases (Agha and Quesada, 2014). In this way, it seems particularly crucial to consider the potential threatening of the ecosystems for all cyanobacterial blooms, even those which don’t produce necessarily MCs. In order to better apprehend the severity of their potential biological effects, the monitoring by MS/MS technologies of phytoplankton biomasses will help to discover new secondary metabolites and variants of known cyanotoxins. This could help, in combination with metabolomics studies on other biological compartments (*e.g.* on zooplankton or fish organisms), to better disentangle the specific and/or synergic metabolic effects of those main metabolite types and thus to bring new understanding of their chemical diversity and respective biological activities.

In addition to the production of toxic secondary metabolites, changes in physico-chemical parameters of the water-bodies have also been reported during cyanobacterial bloom episodes. Massive decrease of dissolved O_2_ concentrations may occur during important bloom senescence events, due to the bacterial degradation of cyanobacterial cell biomass, inducing sometimes spectacular massive death of various fish species, also described as fish-kill phenomenon (Paerl and Paul, 2011). During our study, no noticeable depletion of the dissolved O_2_ concentrations of neither surface nor bottom water or any fish-kill have been observed, indicating that the high cyanobacterial biomasses (observed in V, VS, T and F) were not in senescence. Furthermore, O_2_ concentrations does not appear to significantly drive the liver metabolic differences observed in the fish species studied, suggesting that no metabolic disturbances linked to O_2_ concentration variation were observed here. Among the environmental factors measured on the studied lakes that are noticeably correlated with metabolic differences in fish livers based on rCCA analyses, the pH exhibits wide variations between sampled aquatic ecosystems. Indeed, it appears that lakes dominated by cyanobacteria show higher pH values (above 9), excepted in Grand Fontenay (F) lake where the value is similar (between 7.1 and 8.3) to those found in lakes where cyanobacteria are in minority. Such increase in water pH is an already reported phenomenon directly related with cyanobacteria photosynthesis process that removes carbon dioxide from the water and increases hydroxide ion concentration (Lopez-Archilla et al., 2004). In our study, it is not possible to conclude whether the elevated pH observed in cyanobacterial dominated lakes is a consequence of the photosynthetic activity of cyanobacteria or is due to local geochemical conditions of the respective lake. To better characterize the causality of these elevated pH in further investigation, it should be suitable to monitor pH value before, during and at the end of the bloom.

### Metabolic changes in response to perturbed lakes: cyanobacterial concentrations and pH as environmental drivers of fish liver metabolome

Thanks to NMR and multivariate analyses, our results show that similar metabolic changes are observable in both fish species exposed to high cyanobacterial biomasses (F, VS, T and V lakes). Interestingly, T lake was not characterized by the highest concentration of cyanobacteria observed during our study. However, due to the presence of other pollutants already monitored in past studies in this pound, our observation suggests that additive and/or synergistic effects of multi-pollutants together with cyanobacterial bloom seem to be involved in similar metabolic variations than those of fish from pounds which are the most stressed by cyanobacterial blooms. Nevertheless, as our study was not directly focused on the measurement of the other pollutants present in T lake, further metabolomics studies taking into account the complexity of all those contaminants present together in these ecosystems could provide an even more conclusive observation. However, in our analyses the most constraining factors related to the metabolic changes in fish were the cyanobacterial concentrations and the pH values that exhibit the strongest correlations with the changes of relative concentrations of various metabolites in the two fish species sampled. In this study, it is also very likely that the combination of both high pH values and high cyanobacterial biomasses could lead to the more contrasted metabolome changes in fish. This would be in agreement with the fact that fish from F lake, presenting high cyanobacterial and MCs concentrations but moderate pH value around 7.5, exhibit intermediate metabolomes compared to fish captured in T, V and VS (high cyanobacterial and MCs concentrations and high pH) and fish captured in other lakes (C, CM, M, R), not dominated by cyanobacteria.

Interestingly, MCs concentrations do not seem to be directly correlated with the observed metabolome variations in fish. Our results indicate that MCs may not be in this case the most constraining factor for fish from those natural ecosystems. However, it is also possible that the metabolome differences, correlated to the presence of cyanobacterial cells, could be more generally influenced by the presence of the different bioactive, if not toxic, metabolites present in the various cyanobacterial cells. In addition, as fish sampled in our study were juvenile’s carnivorous species that may mainly feed on zooplankton organisms, as Cladocera or copepods, they may not be exposed by direct ingestion of cyanobacterial cells, avoiding most of the cyanotoxins that remain mainly intracellular during the bloom (Chorus and Bartram, 1999). Nevertheless, they could have been exposed to some amount of the cyanotoxins through those that are extracellularly released in the water or that are contaminating zooplankton organisms feeding directly on cyanobacteria *via* the trophic transfer (Sotton et al., 2014). As it is known that cyanotoxins, such as MCs, are biodiluted along the trophic network (Kozlowsky-Suzuki et al., 2012), it is possible that zooplankton organisms act as ecological filters leading to a decrease in cyanotoxins concentrations before their contact with carnivorous fish. It would be interesting to consider now other fish species feeding on phytoplankton in order to test whether they are indeed exposed to higher cyanotoxin contents, by direct ingestion of cyanobacterial cells, and whether they exhibit similar or even more drastic metabolome alterations than carnivorous ones.

### Perturbations of metabolic pathways in fish: are they stressed?

Metabolomics studies represent a promising step to monitor and potentially early detect environmental stress such as contamination occurring in ecosystems. The specificity of the metabolic signatures could be attempted to be related to every pollutant and would also improve our understanding of their specific physiological effects on fish.

Our results show that common metabolic variations are observed in the two fish species sampled from the cyanobacterial dominated lakes. It appears that all the metabolites highlighted by our relevance network exhibit negative correlations with cyanobacterial content and pH value factors suggesting that lower relative concentrations of these metabolites in both fish species would be found in response to an increase of cyanobacterial content or water pH values. Relative concentrations of metabolites involved in protein and amino acids syntheses, such as homo-L-arginine and L-arginine, were decreased in fish sampled in the cyanobacterial-dominated lakes. Arginine is an essential amino acid implicated in many metabolic functions such as protein, creatine and urea syntheses, the glutamate metabolism and the excretion of insulin and glucagon (Chen et al., 2016). Arginine is then known to increase growth performance in fish, reinforce immune functions and reduce the environmental stress. Thus, a decrease in arginine in fish sampled in the cyanobacterial-dominated lakes could be relied to an up-regulation of the protein synthesis, as it has been previously shown that protein synthesis processes are observed in fish exposed to cyanobacteria or cyanotoxins (Qin et al., 2016; Sotton et al., 2017). Thus, the *de novo* synthesis of detoxification proteins, or the increase of the amino acid catabolism in relation to energy production demand is supposed to undergo the toxic effects of cyanobacteria or other pollutants. Furthermore, these two metabolites could be good candidates as specific biomarkers of a long-term cyanobacterial exposure, as the ANOVA analysis reveals only a significant effect of cyanobacterial biomasses on their concentrations whereas for the other metabolites both cyanobacterial biomasses and pH value had a significant effect on them.

Interestingly, relative concentrations of L-glutamine, a metabolite implicated in alanine/aspartate, amino sugar, D-glutamine/glutamate, nitrogen and purine metabolisms, show also significant lower values in fish sampled in the cyanobacterial-dominated lakes. Glutamine is an important precursor of arginine that is known to repair lipid peroxidation damages due to ROS activity and increase the activity of antioxidant enzymes, at least in humans (Saad, 2012). Furthermore, glutamine is also a precursor of glutathione (GSH) that is implicated in antioxidant reaction and the repair of proteins, and lipid following a ROS production increase. This variation of L-glutamine could be due in part to its transformation in arginine and/or glutathione in order to counteract the ROS over-production, a potential consequence of cyanobacterial metabolites effects.

Relative concentrations of L-palmitoylcarnitine and isovalerylcarnitine were also decreased in fish sampled in cyanobacterial-dominated lakes. These two acylcarnitine metabolites are implicated in cell signaling, fuel and energy storage, fatty acids and lipid metabolisms and in the cell membrane integrity and stability (Reuter and Evans, 2012). Acylcarnitine metabolites are mainly implicated in the mitochondrial β-oxidation of fatty acids in order to produce the acetyl-CoA, entering the Kreb’s cycle, together with NADH and FADH_2_, whose high potential electrons feed the mitochondrial respiratory chain. Disruption of the mitochondrial electron transport chain following an exposure to MCs has already been reported (Zhao et al., 2008). However, the underlying mechanisms of this dysregulation are still unclear. We can here speculate that acylcarnitine metabolites could be related with the mitochondrial function dysregulations potentially induced by cyanotoxins, such as MCs.

Glycerophosphocholine (GPC) variations are also observed in fish from cyanobacterial-dominated lakes. This molecule is a precursor of the phosphatidylcholine that is the major structural phospholipid of cellular membranes. In gills of fish exposed to mercury, the decrease of GPC has been relied to the membrane stabilization/repair processes in order to prevent lipid peroxidation damages (Cappello et al., 2016). It is well known that cyanotoxins, and notably MCs, induce lipid peroxidation damages in cell (Ferrão-Filho and Kozlowsky-Suzuki, 2011). Additionally, high values of water pH could be also implicated in the denaturation of cellular membrane (Zahangir et al., 2015). Our observation suggests that these GPC low contents could be related to cellular repair processes in order to counteract membrane damages due to the lipid peroxidation potentially induced by cyanobacterial metabolites and/or high pH values. Interestingly, as glycerol is implicated in glycerophospholipid metabolism, its low concentration in fish exposed to cyanobacterial-dominated lakes, could reflect its consumption in order to produce new membrane glycerophospholipids required for the repair of the lipid peroxidation damages to membranes.

Furthermore, the relative concentrations of two steroid compounds, the 16a-hydroxyestrone and the androsterone were present in a lesser amount in fish livers from cyanobacterial-dominated lakes. The 16a-hydroxyestrone and the androsterone are biotransformed metabolites of estrogens and androgens respectively, playing in fish critical roles in various reproduction, growth, and developmental processes. Endocrine disruption effects in adult model fish (zebrafish and medaka) following experimental exposures to cyanobacterial blooms or cyanotoxins have already been related in the recent literature (Liu et al., 2018; Sotton et al., 2017). However, our results suggest that similar endocrine disturbance could occur in field where fish are exposed to even longer period of cyanobacteria and cyanotoxins occurrence.

Finally, relative concentrations of the acetylcholine, a neurotransmitter, were present to a lesser amount in fish from cyanobacterial-dominated lakes, being in potential link with neurotoxic effects of cyanobacterial blooms. MCs have already been shown to induce a decrease of acetylcholine concentrations in zebrafish embryo leading to neurodevelopmental disturbances (Wu et al., 2016). The effects of this neurotransmitter dysregulation could indeed be particularly critical in juvenile fish, as they are still in development. Furthermore, it is possible that other cyanobacterial secondary metabolites could dysregulate neuronal systems, as it has already been shown that some cyanobacteria are able to produce specific compounds with neurotoxic effects (Aráoz et al., 2010). In this way, it seems particularly important in field studies to perform LC-MS/MS analyses of cyanobacterial bloom biomasses in order to better characterize the diversity of cyanobacterial compounds potentially impacting the biology of aquatic organisms.

### Conclusions and horizons

Our study demonstrates that cyanobacterial blooms can induce locally metabolic variations in relation to environmental stress response of fish. Such metabolomic analyses support already known results but give also new perspectives for the characterization of the cyanobacteria-fish interactions. In this way, it could be interesting to do further studies, thanks to longitudinal samplings, on the changes of fish metabolomes during bloom episodes, which could help us to better understand and characterize the main disturbing environmental factors during these anthropo/ecological phenomenons. Also other species feeding on phytoplankton organisms should be sampled in order to highlight whether higher metabolome changes could be observable due to the fact that they directly feed on cyanobacterial cells and thus are in direct contact with high concentrations of various potentially toxic secondary metabolites. Although NMR approach represents a useful tool to classify, monitor and potentially detect early disturbances related to cyanobacteria and more generally to environmental stresses occurring in ecosystems, the parallel investigation by LC-MS/MS could be particularly useful in order to give an even better strength of the analyses. Indeed, it could gain at confirming the molecular identification and quantification and bring additional information especially concerning the different metabolites present only in limited amounts. Furthermore, complementary Omics analyses as the study of the proteome and the transcriptome in addition to metabolome information will help us to better characterize potential biomarkers of environmental stresses, and to depict the mechanism of the physiological effects of environmental stressors on fish populations.

## Acknowledgments

The NMR and the MS spectra were respectively acquired at the Plateau technique de Résonance Magnétique Nucléaire and the Plateau technique de spectrométrie de masse bio-organique, UMR 7245 CNRS/MNHN Molécules de Communication et d’Adaptation des Micro-organismes, Muséum National d’Histoire Naturelle, Paris, France.

## Author Contribution

B.S., A.P., S.L.M., Q.Q., A.C., C.B. and B.M. conceived the sampling strategy. B.S., S.L.M., Q.Q., C.D., C.B., and B.M. conducted the sampling. B.S., A.P., S.L.M., A.B., A.C., V.P. and B.M. performed the analyses and analyzed the results. All authors reviewed the manuscript.

## Author information

### Corresponding author

UMR 7245 MNHN/CNRS Molécules de communication et adaptation des micro-organismes, Muséum National d’Histoire Naturelle, 12 rue Buffon, F-75231 Paris cedex 05, France.

Tél: (+33) 140 793 179; &

